# Unfeasible expectations: the sensitivity of structural stability measures favours simpler metrics for empirical questions

**DOI:** 10.1101/2025.04.24.650476

**Authors:** J. Christopher D. Terry

**Affiliations:** Department of Biology, University of Oxford, Life and Mind Building, South Parks Road, Oxford OX1 3EL

**Keywords:** coexistence, structural stability, ecological networks, *Drosophila*, ecological forecasting, empirical uncertainty

## Abstract

Understanding what determines community assembly and disassembly in a changing environment is a core challenge for ecology. Recently a family of structural stability approaches that determine the range of intrinsic growth rates compatible with system feasibility have been gaining popularity as a measure of how likely a community is able to persist in fluctuating conditions. This offers a theoretical basis for understanding and predicting the assembly and stability of complex multi-species communities from only interaction network structures. However, here I show that the high sensitivity of calculations of the feasibility domain, coupled with empirical uncertainties inherent in estimated interaction strength, are likely to preclude the approach’s reliable application to empirical settings as a metric to compare the stability of different communities. Across four reanalyses of previous empirical demonstrations of the structural stability approach, more parsimonious measures based on species connectance provide better explanations for patterns of community assembly or differences in stability. Calculation of structural stability metrics therefore appears to lose, rather than synthesise, information embedded in empirical interaction matrices. This success of simpler measures is good news for the purposes of prediction and emphasises the value of multiple-competing hypotheses in validation tests to demonstrate value-added associated with new approaches.

## Introduction

Understanding the conditions and processes that determine the coexistence or otherwise of species is a central goal of ecology, but one that is hampered by the substantial, and inevitable, deficits in our empirical knowledge of species responses and interactions. Simplifying theoretical constructs are necessary to allow tractable analysis of complex natural ecosystems. A perennial challenge is how to best navigate the balance between pressures for increasingly detailed mathematical frameworks and metrics with the limitations of available empirical knowledge.

Recently the structural stability approach (Rohr *et al*. 2014; Saavedra *et al*. 2017) has been gaining popularity as a tool in the quest for understanding and predicting both coexistence and assembly patterns of ecosystems. At its core, it takes an assumed matrix describing interactions between species under the framework of Lotka-Volterra models to calculate the diversity of population intrinsic growth rates across the community that result in feasibility - all species having positive equilibrium biomasses. A relatively large feasibility domain implies that interspecific interactions impose relatively weak constraints as species can coexist at a larger number of intrinsic growth rates combinations. As such, a community that has a larger feasibility domain is seen as relatively more stable. By contrast, a smaller feasibility domain implies that constraints are more rigid and under inevitable fluctuations the community is more likely to collapse. This is frequently interpreted probabilistically as capturing the likelihood that a random combination of population intrinsic growth rates result in a feasible community (Deng *et al*. 2024b; Song *et al*. 2020b).

Feasibility is often touted as a more ecologically relevant response variable than other options such as local asymptotic stability, and has been proffered as a means to understand the impact of external changes on communities given only knowledge of an interaction matrix (Allen-Perkins *et al*. 2023; Godoy 2019; Godoy *et al*. 2024; Saavedra *et al*. 2020). The approach can unite multiple strands of ecological thought (Godoy *et al*. 2018) and facilitates detailed analytic treatment of communities larger than the pairwise analyses possible under, say, classic Modern Coexistence Theory (Barabás *et al*. 2018; Chesson 2000). It allows the same framework to be used for both pairwise analyses (Song *et al*. 2020a) through to large communities (Grilli *et al*. 2017).

However, applying the standard structural approach to an empirical system is ultimately predicated on a given interaction matrix being a suitably accurate description of the interrelationships within the ecological community (Saavedra *et al*. 2020). Unfortunately, although the qualitative topology of the direct interactions between species may be directly observed or predicted from traits, quantitative interaction matrices are notoriously challenging and effortful to even approximately parameterise (Dormann 2023; Wootton & Emmerson 2005). While tools for predicting interaction strengths from traits have shown promise (Berlow *et al*. 2009; Freckleton & Watkinson 2001), accuracy still has a low ceiling. Furthermore, interactions frequently vary substantially through time as they are highly dependent on the environmental context (Chamberlain *et al*. 2014; Tylianakis *et al*. 2008), biotic conditions (Peacor & Werner 2004) and life-history stage (de Roos 2021), making extrapolations from controlled conditions a considerable challenge.

Strong sensitivity to the quantitative details of interaction matrices has been a hallmark of stability results using other metrics due to the rapid indeterminacy arising from chains of indirect effects (Novak *et al*. 2011; Yodzis 1988). Under the standard structural approach, the uncertainty and probabilistic interpretation are pushed onto the intrinsic growth rates, despite the observation that in many communities estimates of interaction coefficients are much more uncertain than intrinsic growth rates (e.g. Terry 2024). Given these challenges, it is therefore very notable that the structural stability approach has had multiple high-profile recent successes in describing the assembly of empirical communities (Bartomeus *et al*. 2021; Deng *et al*. 2024a; Domínguez-Garcia *et al*. 2024; García-Callejas *et al*. 2023; Medeiros *et al*. 2021; Saavedra *et al*. 2016; Song *et al*. 2018a; Tabi *et al*. 2020) despite often having access to only relatively crude approximations of interaction matrices, for example those generated from qualitative resource-use overlap approximations.

Ecological theory has many potential valid objectives (Grainger *et al*. 2021; Marquet *et al*. 2014; Shaw *et al*. 2024) including deeper understanding of potential mechanisms and improved predictions of observed dynamics. For the objective of assessing the stability of different communities, identifying a correspondence with data can only be a starting point of validating the utility of an approach. Ideally, a useful abstraction or theory should capture something additional and distinct to more parsimonious approaches.

Here, I first demonstrate the sensitivity of structural stability metrics is such that even small empirical uncertainties in the input interaction matrix can be expected to overwhelm any real shifts in a communities’ feasibility domain and examine the challenges of propagating uncertainties to explore the consequences of focussing the uncertainty onto species intrinsic growth rates. I then reanalyse four previous results to identify alternative simpler explanations for identified correspondences between inferred structural stability and observed community dynamics. I argue that, since apparent successes of structural stability are frequently exceeded by (or at the very least correlated with) simpler properties of the community, it may well be more profitable to focus on these simpler drivers when assessing the stability of empirical communities. Ultimately this is a good news story, demonstrating the valuable predictive information contained within even the relatively crude interaction networks available.

### Calculation of a feasibility domain

This section briefly summarises approaches to calculating the size of a feasibility domain (′Ω’, or when scaled per species ‘Ω’) given an interaction matrix **A**. To avoid simply repeating the many previous thorough treatments (e.g. Saavedra *et al*. 2017; Song *et al*. 2018b; Svirezhev & Logofet 1983), here the intuition is emphasised to keep the technical mathematical language to a minimum.

The population dynamics of each species (*N*_*i*_) are assumed to follow Lotka-Volterra dynamics: 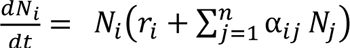, with intrinsic growth rates *r*_*i*_ and competition coefficients from species *j* on to species *i*. However, it can also be applied to other similar models, such as alternative formulations of Lotka-Volterra (e.g. Tabi *et al*. 2020), nonlinear functional responses (Cenci & Saavedra 2018) or seed-bank models of plant dynamics (Song *et al*. 2021). This model can be written in matrix form and solved for equilibrium, **0 = *r* + **AN**^∗^. The equilibrium densities for a given community specified by *r* and **A**, is then *N*^∗^ = −**A**^−1^ ⋅ *r*. The fundamental question for a feasibility analysis is then, given only information on **A**, what values of *r* would lead to all values of *N*^∗^ being greater than zero?

The possible space of *r*_*i*_ values to search over is made easier by the observation that scaling the vector of *r* values wouldn’t change the sign of *N*^∗^. This means that each ratio of *r* values need only be tried once, which can be achieved by normalising the set of *r*’s in some way. Approaches include the ‘L1 norm’ (*r*_1_ + ⋯ + *r*_n_ = 1) or more commonly the ‘L2 norm’ 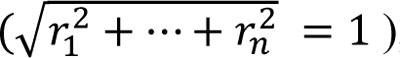, which embeds the assumption that each ratio is equally likely and has the visually attractive property of forming a sphere in three dimensions (e.g. Fig. 1b).

**Figure 1.**
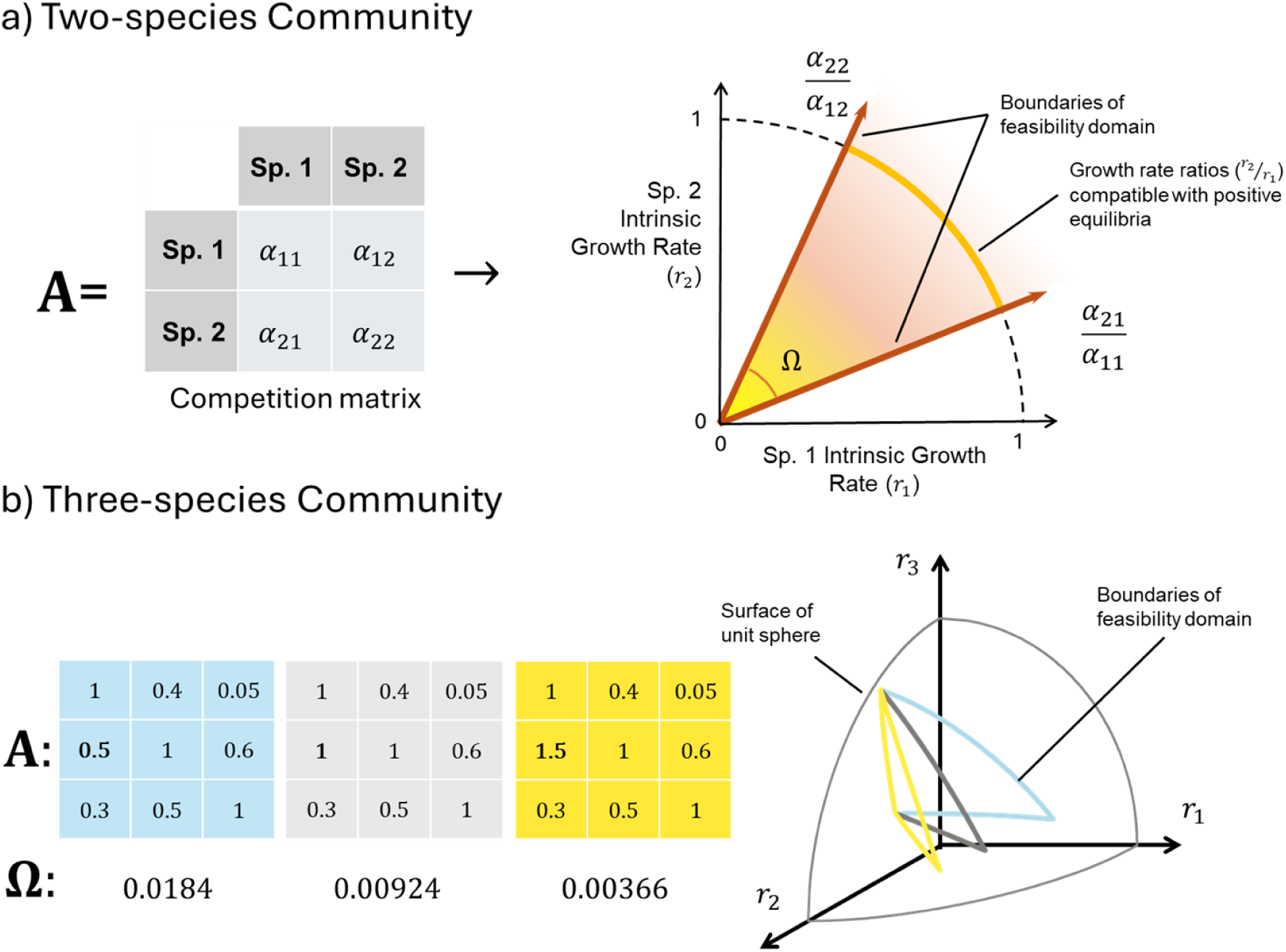
Illustration of calculation of feasibility domains for small communities. The objective is to calculate the proportion of combinations of intrinsic growth rates (*r*) that are compatible with all species having a positive equilibrium (‘feasibility’), given an interaction matrix **A** and assuming Lotka-Volterra dynamics. a) With two species, the boundary of the feasibility domain can be determined analytically, and the proportion Ω of feasible combinations of *r* is the angle between these boundaries. b) With three species, this angle becomes the triangular surface area on a sphere. Here three domains corresponding to the three matrices are shown in three colours. Because all interactions are of the same sign, only one sector of the sphere is relevant. Note the large response in both the size Ω and location of the domain to a changing only a single element (α_21_, in bold) by ±50%. Beyond three species, the domain is no longer easily visualised, but the same principle applies.

With the Lotka-Volterra model, the possible *r* values that lead to feasibility make up a continuous ‘domain’, within definable boundaries. This feasibility domain can be summarised by both *which* vectors lie in the feasibility domain, and proportionally *how many* vectors. For two species, these boundaries, and hence the size of the feasibility domain Ω can be calculated directly (Fig. 1a). For more diverse communities, the calculation becomes a complex integral across each of the species. Fortunately, this can be computed through a quasi-Monte Carlo approach for fairly large communities (Genz & Bretz 2002; Saavedra *et al*. 2016) and these functions are easily accessible through public R packages such as *feasibilityR* (Saavedra *et al*. 2023). Effectively, this samples the proportion of a large number of trialled *r*-value combinations spanning the full range of possibilities that lead to feasibility. This proportion is denoted Ω (sometimes Ξ) and captures the *size* (although not the *location* or *shape*) of the feasibility domain. There also exist analytic approximations for large communities where the interaction strengths are identically independently distributed (Grilli *et al*. 2017).

Since with increasing species diversity, it tends to get harder to identify combinations of *r* that lead to feasibility, it is common to instead report this value scaled by the number of species n, ω = Ω^1/n^. As such, ω can be interpreted as the average probability that a species will have a positive equilibrium value with a randomly drawn *r*, making the chance of the whole community being feasible ω^n^ = Ω. The size of the ‘possible’ domain can be restricted to increase the biological meaningfulness, for example only considering positive *r* values in a competitive community, or negative values in a mutualistic one. These restrictions can scale the bounds of maximum feasibility.

It is worth noting that *feasibility* does not imply *stability* by default. Without even local stability feasibility on its own can’t be taken to imply coexistence. However, feasibility is often associated with local stability in sampled systems (Case & Casten 1979; Roberts 1974; Song & Saavedra 2018; Stone 2018). Local asymptotic stability is much more likely with relatively strong intraspecific interaction strengths (Barabás *et al*. 2016, 2017) and can be guaranteed under certain conditions – for example the negative definite matrices studied by Grilli *et al*. (2017).

### Sensitivity of Calculation of Structural Stability

Uncertainty in the estimation of feasibility domains enters in multiple ways with varying degrees of tractability. Numerical instabilities in the calculation of very small feasibility domains (generally when n > 10) can be addressed by brute-force repeated calculation. However, underlying uncertainty in **A** is a more fundamental challenge. ‘Probabilistic’ interpretations of ω as the chance that a species will be able to persist given its interactions (Song *et al*. 2020b) must consider the contributions of uncertainty in interaction strength and the space of possible r-vectors.

Sensitivity analyses of the impact of perturbations to **A** are simultaneously a test of the impact of misidentification of measured interaction strengths and an analysis of a systems response to environmental changes affecting interactions. These are fundamentally different interpretations, but in an empirical application of the framework it is essential to separate the first from the second. While responsiveness to small changes is potentially an advantage in a tool, this is only the case if the underlying inputs are measurable with sufficient precision, something that is unfortunately not the case for ecological interaction networks (Wootton & Emmerson 2005). Distilling the complexities of real ecological communities into an interaction matrix is an essential but challenging process requiring significant assumptions about the state and dynamics of the system (Laska & Wootton 1998; Novak *et al*. 2016). The underlying data to generate these matrices is commonly the frequency of observed interactions between species which must be converted into interaction coefficients with an assumed population dynamics model (e.g. Rohr *et al*. 2014). While it is long established that empirical interaction matrices can capture important and interesting features of ecological communities, they cannot be expected to be highly precise, and are dependent to their assumptions (e.g. Cervantes-Loreto *et al*. 2023).

The numerical calculation of the feasibility domain is based on a matrix inversion ((*A*^*T*^A)^−1^). While inverting the resultant symmetric matrix is somewhat more robust than an arbitrary matrix, errors in elements of **A** will almost always magnify, rather than be averaged out through this inversion process (Barabás & Allesina 2015). This is a longstanding issue that applies to any metric that relies on matrix inversion (Yodzis 1988) and the exact sensitivity is dependent on the structure of the matrix being inverted (see e.g. Iles & Novak 2016; Pomerantz 1981). Numerical examples can get a sense of the problem (Fig. 2); however, understanding if the error-driven variability identifiable in such artificial tests is problematic not straightforward because it depends on the relative balance of ‘real’ and ‘uncertainty-driven’ differences in feasibility and the number of communities being compared. Although ω can take any value between 0 and 1, the span of reported ω values for different communities within particular empirical studies are often much narrower (e.g. approximately 0.45-0.55 (Domínguez-Garcia *et al*. 2024), 0.76 – 0.79 (Song *et al*. 2018a)). It is therefore critical to explore the role of uncertainty in **A** for empirical systems and to compare the performance relative to other metrics.

**Figure 2.**
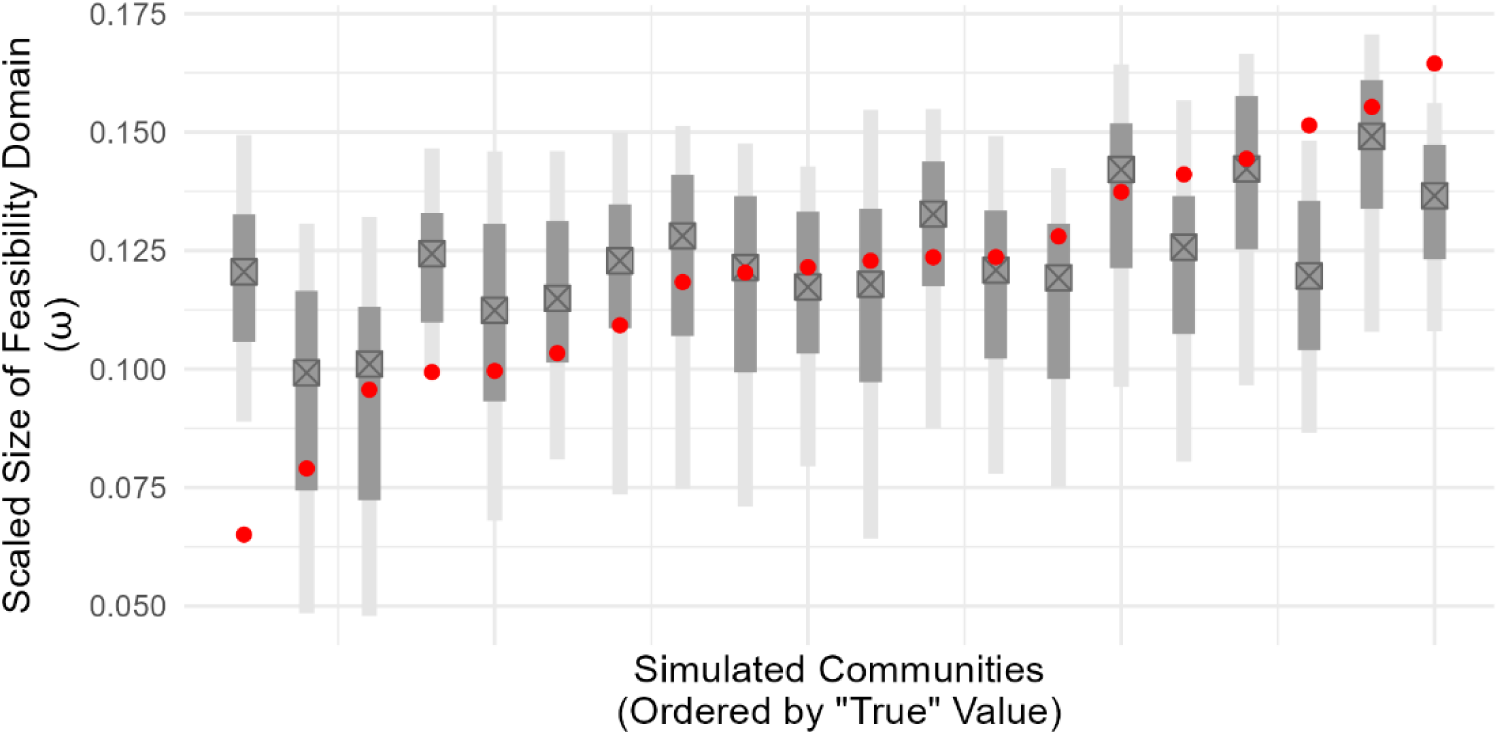
Demonstration of sensitivity of feasibility domain to error in elements of. **A**. Twenty artificial communities of 15 competing species were generated with a ‘true’ interaction matrix **A**^*True*^with connectance 0.5 parametrised with draws from a negative half-Normal distribution (mean 0, sd = 2). The diagonal entries were fixed to −1. Scaled feasibility domain size (ω) of **A**^*True*^ranged over approximately 0.1 (red dots). Grey lines show median, 66% and 95% intervals across 100 separate applications of measurement error added by perturbing all non-zero elements of **A**^*True*^by ±20%. This represents moderate interaction strength uncertainty while still retaining the overall topology nearly perfectly (e.g. average rank correlation of the non-zero entries between original and perturbed **A** was 0.96). This error was sufficient to largely obfuscate the underlying ‘true’ differences in feasibility domain. True values frequently fell outside the range estimated under error and across the draws a Spearman’s Rank test between the ‘observed’ and ‘true’ measures could only identify a significant (p<0.05, n= 20) correlation in less than half (41%) of cases. These results are however strongly dependent on the structure of the original interaction matrices and the errors applied to them.

### Empirical two-species example incorporating empirical uncertainty

Interpretation of ‘virtual ecologist’ power analyses are inevitably limited by the need to specify both the level of real variation to be discerned and the level of empirical noise. To give a more concrete example of the impact of parameter uncertainty on the calculation of feasibility domains, Fig.3. illustrates an example where interaction parameters and uncertainty was directly estimated across a range of temperatures. The data come from a comparatively well-characterised two-species *Drosophila* mesocosm system described in Terry (2025). With high replication (thousands of observed transitions across the two species) and a highly controlled environment, the precision of these parameter values is likely to represent a ‘best-case’ scenario. The two species have distinct thermal preferences and model selection identified shifts in the interaction coefficients with temperature. The fitted model had a Beverton-Holt form from which structural stability can be calculated (Song *et al*. 2021) and posterior estimates for the parameters are given in Supplementary Information 2. Since in this case the intrinsic growth rates of each population at each temperature are known this is arguably not a scenario where structural stability would be considered a core tool (although see Song *et al*. 2021). However, the additional knowledge allows investigation of the application of the approach to cases where only interaction matrices are available. A pairwise case study allows easier visualisation of the challenges that are likely to be more problematic at higher dimensions.

**Figure 3.**
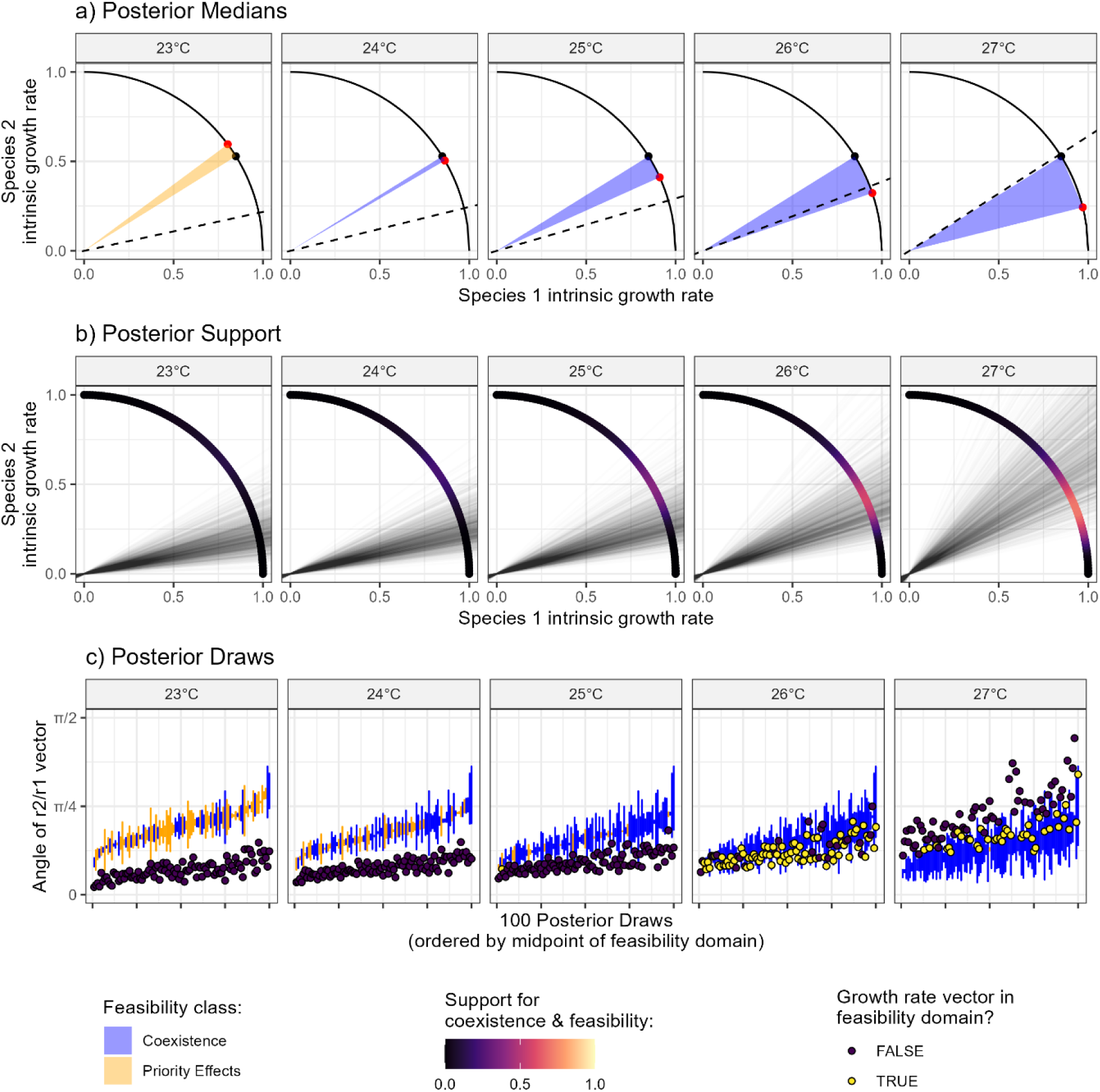
Feasibility domains of the competition between *D. pandora* and *D. pallidifrons* at five temperatures using fitted parameter posterior distributions. **a)** Feasibility domains using posterior median parameter values. Dashed line shows median fitted intrinsic growth rate ratio. b) Support for each point on the unit sphere being part of the feasibility domain across the full posterior given by colour. Note that most points have low support for being contained in the feasibility domain. The fan of lines illustrates the posterior distribution of growth-rate vectors. c) Fitted growth rates are not independent of fitted competition terms. To illustrate this feasibility domains and growth rate vectors of 100 posterior draws are shown for each temperature.

A direct structural analysis making use of the posterior medians (Fig. 3a) would identify a marked rise in the size of the feasibility domain from 24°C with temperature. Note that there was only statistical support for identifying an impact of temperature on the α_11_ and α_21_ terms, hence only one of the feasibility domain boundaries responds to temperature. However, when considering the full posterior distribution of the parameter values, several challenges become clear:

Firstly, the *overlap* of feasibility domains from different draws of the posterior is at best moderate. In Fig. 3b, it can be seen that the support that any given community intrinsic growth rate vector (*r*_2_/*r*_1_) is part of the identified coexistence feasibility domain is very rarely above half. Indeed, the overlap in feasibility domains varies more due to experimental imprecision than by an environmental change (+4°C) that was sufficient to reverse the outcome of competition. This poses a severe challenge to approaches that hope to measure the impact of environmentally driven changes from the overlap in feasibility domains (Godoy *et al*. 2024; Song *et al*. 2018b) as any such shift will likely be overwhelmed by even moderate empirical uncertainty. It would also be expected that the overlap of higher-dimensional feasibility domains between posterior draws would reduce faster in more diverse communities.

Second, the status of the feasible equilibria within the domain, i.e. whether stable coexistence or priority effects (Song *et al*. 2020a) are expected, is not necessarily consistent under parameter imprecision. Where the boundaries of the feasibility domain ‘cross’ (α_21_/α_11_ > α_22_/α_12_, making niche overlap 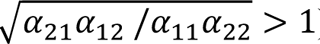), the interpretation of “feasibility” changes from stable to unstable equilibria. Different expressions to calculate the size of the feasibility domain implicitly deal with this challenge in different ways (Supplementary Information 1), but the widely used multivariate Gaussian sampling approach to measure Ω does not distinguish between the classes of “stability”. A further consequence is that the size of the feasibility domain calculated from the median parameter values (Fig. 3a) is identified as shrinking between 23°C and 24°C (to a very small value) while examining individual draws from the full posterior (Fig. 3c) shows a consistent increase – the difference comes from averaging over a mixture of both stable and unstable feasibility domains.

As a distinct issue, in this system it is known that at higher temperatures that *D. pandora* consistently and rapidly excludes *D. pallidifrons* (Terry 2025). Despite these challenges around quantifying the feasibility domain’s location and size, it is nonetheless clearly predicted that the feasibility domain would enlarge with temperature above 24°C. Without information of the intrinsic growth rates (the standard application of the structural approach), this would be interpreted as implying that coexistence of the two species was increasingly likely, the opposite of what is experimentally observed. Although there is increasing feasible space in the *r*_2_/*r*_1_ simplex, at higher temperatures the observed *r*_2_/*r*_1_vector moves more rapidly anti-clockwise through and away from this region (Fig. 3a). While this is clearly just a single example, the relative size of the domain can be less important than its location or, in more diverse communities, its shape relative to the response of growth rate vectors.

### Re-evaluating Case studies & Alternative Explanations for Success

Recently there have been an increasing interest in extending the study of feasibility domains from a purely theoretical pursuit to empirical settings (Allen-Perkins *et al*. 2023; Saavedra *et al*. 2020). Multiple studies have identified relationships between the size of the feasibility domain and various measures of community assembly and stability, without the sample sizes that might be expected to be needed given the expectation of noise identified above. This is especially surprising given that many analyses are ultimately based on binary bipartite interaction matrices, from which are calculated (non-binary) effective interaction matrix based on inferred resource use overlap (Rohr *et al*. 2014, Fig. S1). While this approach has a long pedigree and makes best use of the available data (Godoy *et al*. 2018), it cannot be reasonably expected to generate **A**-matrices with the high levels of precision to give a dependable estimate of ω.

What then can explain the success? There are enough achievements to rule out luck or other bad faith explanations. Here I reanalyse four distinct studies that use empirical data of natural communities, in each case following as close as possible to the original objectives and approaches but testing if structural stability metrics are adding value compared to simpler species-level predictors that may offer plausible parsimonious explanations for the apparent success of calculated feasibility metrics as a correlate of observed community dynamics. These case studies were chosen based on having clearly tested links between structural stability and observed dynamics, openly accessible data and as they had been repeatedly cited as providing an empirical justification for the use of structural stability metrics. R Markdown documents detailing all steps are available in the code repository.

### Case Study 1 – Persistence of pollinator communities across sites

In a large, 6-year, study within southern Spain, Domínguez-Garcia *et al*. (2024) found a strong (ρ = 0.85) positive correlation between the inferred structural stability ω of pollinator communities and average species persistence (as measured by the average proportion of years pollinators are observed in) across 12 sites. Here ω values were calculated based on effective interaction matrices for each site derived from plant resource overlap between the pollinators (Fig. S1). Site ω values are almost perfectly correlated (ρ = 0.96, Fig. 4a) with the nestedness of each site’s plant-pollinator network (calculated by the binary method of Bastolla *et al*. (2009) following the original paper). As might be expected given such a tight correlation, nestedness is indistinguishable from ω as a predictor of pollinator average persistence (Fig. 4b,c, 95% confidence intervals of Spearman’s ρ_ω_= 0.52: 0.978, ρ_nestedness_ = 0.295:0.986), making it hard to distinguish with confidence the underlying driver. However, the original study also identifies moderate species-level correlations within each site between species-specific feasibilities (ω_i_) and species persistence (ρ’s between 0.37 and 0.66 for the 12 sites), which allows alternatives to be tested. Simply using the observed degree of each species within each site’s network gives substantially higher within-site correlations with the observed persistence (ρ’s between 0.706 and 0.93 for the 12 sites, Fig. 4d). Although the original study conducts network randomisations to identify that the specific network structure determines the calculated feasibility values, the process of calculating feasibility does not bring an identifiable gain of information at the community scale compared to the more empirically-robust nestedness metric (Nielsen & Bascompte 2007), and loses considerable predictive power at the species-level.

**Figure 4.**
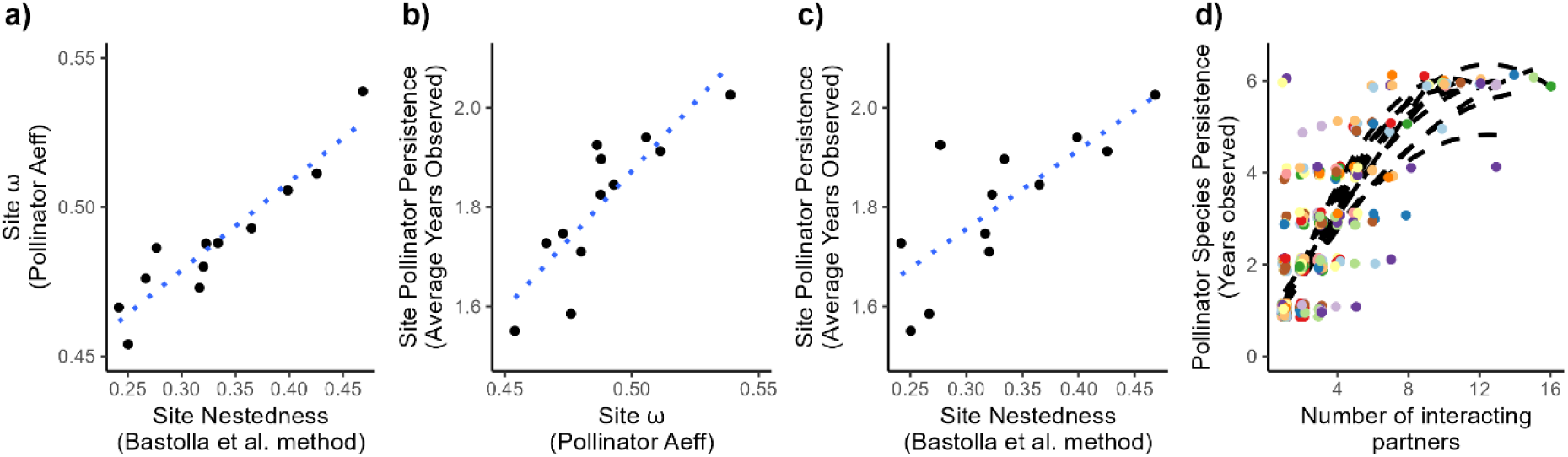
Reanalysis comparing binary nestedness or connectance with Ω as a predictor of pollinator persistence. In a-c, site level metrics are plotted, while in d points represent individual species coloured by site. a-c) Tight correlation between nestedness, the size of the feasibility domain and average persistence of pollinators as calculated by average years (out of 6) each species was seen in. d) At each site, how well does the number of interaction partners of each species predict the species-level persistence, calculated at the number of years (out of 6) that the pollinator was seen in. In all cases Spearman’s rank correlations were used, fitted lines are shown to aid interpreation: a-c are linear model fits, while those in d) are quadratic lines of best fit and the points are jittered to reduce overlap.

### Case Study 2 – Long-term Assembly of a Plant-Herbivore Community

Song *et al*. (2018a) examined how the assembly order of plant species into a large central European region (Baden-Württemberg, 35 751 km^2^) affected herbivore community assembly, taking advantage of a large plant-insect interaction database (Altermatt & Pearse 2011) and estimated arrival times from archaeobotanical and historical records. The observed binary interaction matrix was used to infer an effective competition matrix between insect herbivores that share the food plants present in the region, in each time slice between arrivals. After each set of plant arrivals, the herbivore competition matrix was recalculated and its ω calculated. They found a strong positive correlation between ω and time (Fig. 5a), and showed that this pattern was not simply an artefact of increasing diversity as it was reversed when the order of plant arrival was randomised. Moreover, the correlation was only seen in non-ornamental plants, while no such correlation was observed when focussing only on ornamental plants, suggesting a role for competition in community assembly. This was interpreted as the herbivore community being driven towards network structures with high overlap of host plant use during the early assembly stages, but this overlap diluted as the community developed.

**Figure 5.**
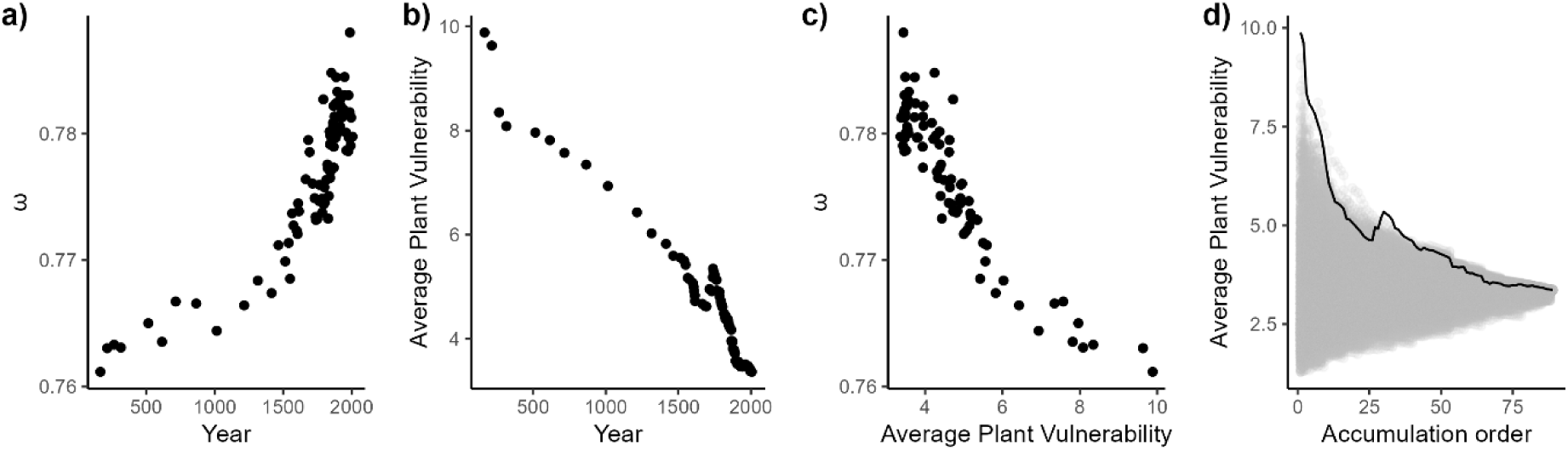
Reanalysis of relationship between plant arrival order into Baden-Württemberg and interactions with a lepidopteran herbivore guild. **a)** Original result of an increase in ω of the herbivore community through time. b) Strong and consistent decline in average links per plant (vulnerability) in the herbivore-plant network. c) Strong negative correlation between connectance and ω. d) Observed trend in average plant vulnerability through time (black line) is markedly non-random when compared to 1000 randomised arrival orders (grey lines).

However, reanalysing the data shows that a stronger trend can be identified from the observation that early arriving plant species tend to have more herbivores consuming them. Average vulnerability (number of herbivores linked to each plant) of the plants in the network in each time slice is highly correlated with time (Fig. 5b). Community average vulnerability and ω are highly correlated (Fig. 5c), but average vulnerability is more strongly negatively correlated with year (ρ = −0.932 [-0.955:-0.898]) than ω is correlated with year (ρ = 0.877 [0.818:0.917]), following the original paper in assuming independence of the time slices. The lack of a pattern with the ornamental plants is again retrieved (Fig.S2). Comparing the observed trajectory of average plant vulnerability though time with 1000x randomised arrival orders highlights the predictive power of this simple signal (Fig 5d). Since there is no comparable data on the arrival of the insect herbivores, the drivers of increased vulnerability of earlier arriving plant species is indeterminable, at least from this dataset. That said, the reanalysis suggests that the detected signal in feasibility of the herbivore community is readily explained by greater accumulation of enemies on earlier arriving plant species without needing to invoke community-level competition explanations at the regional scale.

### Case Study 3 - Species level feasibility domain metrics as a predictor of population dynamics

Allen-Perkins *et al*. (2023) analyse data from Mediterranean annual plants to test whether species-specific metrics of the shape of the feasibility domain have the capacity to predict annual changes in the abundance of seven species across 5 year-year transitions. Specifically, they test the predictive capacity of a species’ ‘exclusion ratio’, a measure that compares the inferred size and shape of the feasibility domain to a null scenario where all species interact identically (Desallais *et al*. 2025; García-Callejas *et al*. 2023). The larger a species’ exclusion ratio, the greater the relative probability that it will be the first species excluded under a random environmental change that perturbs intrinsic growth rates. This therefore allows a species-level assessment of vulnerability within the wider structural stability framework.

The interaction matrices were calculated for each of seven years from independent data from fitted population models (García-Callejas *et al*. 2021). One year was excluded, generating 5 usable transitions. The original authors identified a negative relationship between the exclusion ratio of a species and its population growth rate (log_10_(N_t+1_/N_t_ + 1)), although this is not statistically significant (Fig. 6a). Across all observations Spearman’s ρ was −0.096 (p-value = 0.58, n =35), following the original authors in treating the 5 observed transitions of 7 species as independent.

**Figure 6.**
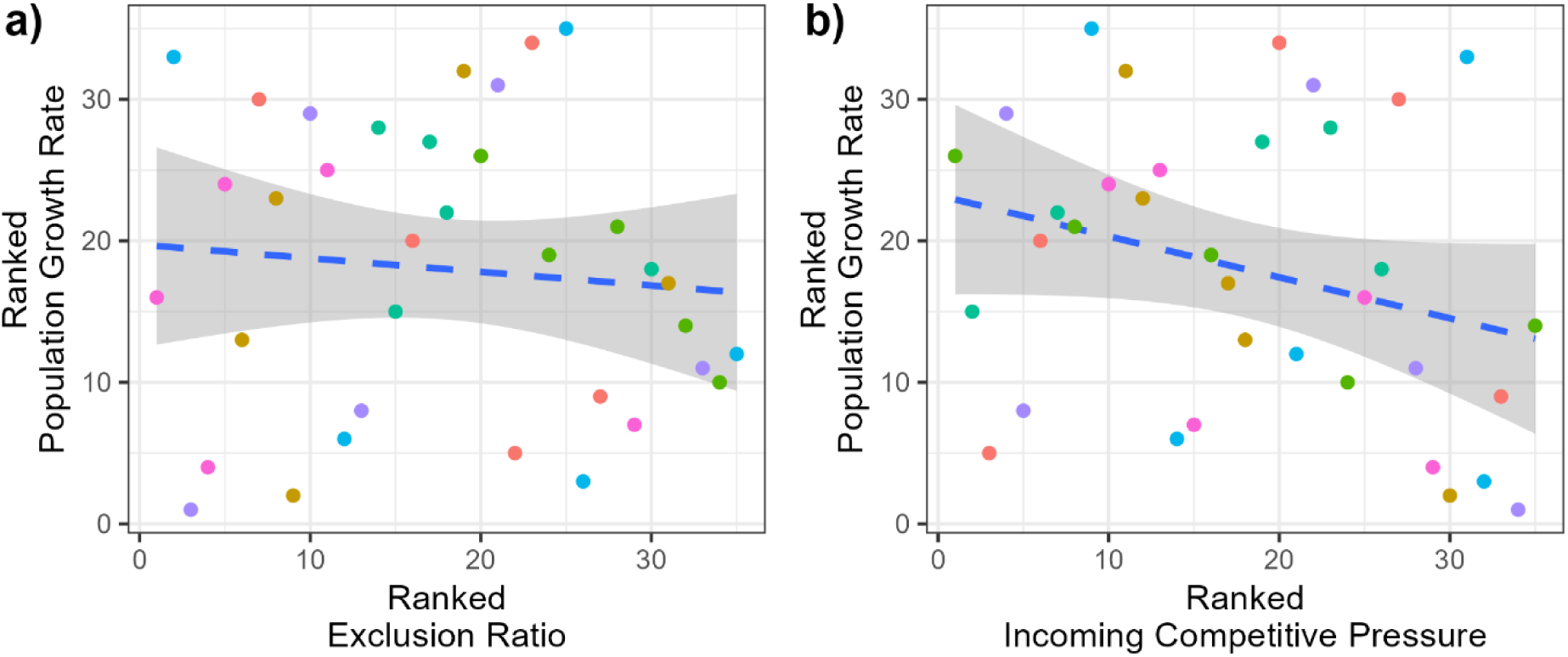
Comparison of explanatory power of species-specific structural stability-based metric to simple competitive pressure total for Mediterranean grassland plant dynamics. **a)** Replication of original result identifying link between species-level exclusion ratio feasibility-based metric that calculated the relative distance to the edge of feasibility domain for each species and observed annual population growth rate. b) Reanalysis using a (crude) estimate of incoming competitive pressure shows a stronger, although still insignificant, negative correlation. In both plots colours indicate different species and the line is a linear model (with 95% confidence intervals) fit through the ranks included to help visualisation.

Given the diffuse competition networks, there is not a direct equivalent of linkage degree to test as a basic predictor. However, a simple, if somewhat naïve, approach simply rowwise sums the interspecific elements (Σ_*j*_α_*ij*_, for all *j* ≠ *i*) of the interaction matrices to give an estimate of the competitive pressure felt by each species. Despite not considering the density of competitors this is able to give a markedly improved, although still statistically insignificant, correlation with observed abundance changes (Spearman’s ρ= −0.299, p-value = 0.091, Fig.6b).

### Case study 4 – Observed combinations of herbivores through time across sites

Medeiros *et al*. (2021) studied the community assembly of caterpillar herbivores across tropical dry forest sites at three successional stages (Boege *et al*. 2019). For each of 88 sampled communities an effective competition matrix **A** between herbivores was constructed based on species occurrences within a plot and a binary meta-network of plant-caterpillar interactions. The Ω of this matrix was calculated and compared to a distribution of Ω(**A**) calculated from equally sized herbivore communities drawn at random from the regional species pool. They found strong support that the observed communities had a larger Ω than would be expected from random draws, and inferred that observed communities are those more likely to persist in the face of changing environmental conditions (Fig. 7a). However, a much stronger signal is seen with a considerably simpler connectance based approach (Fig. 7b). Observed plant-herbivore connectance was much higher than random, and was much more statistically significant than the feasibility domain signal. In fact, in half of the communities, the observed community had a higher connectance than all 10’000 randomly drawn communities. Patterns in community assembly can in this case be directly explained simply by the over-representation of generalist species at each site.

**Figure 7.**
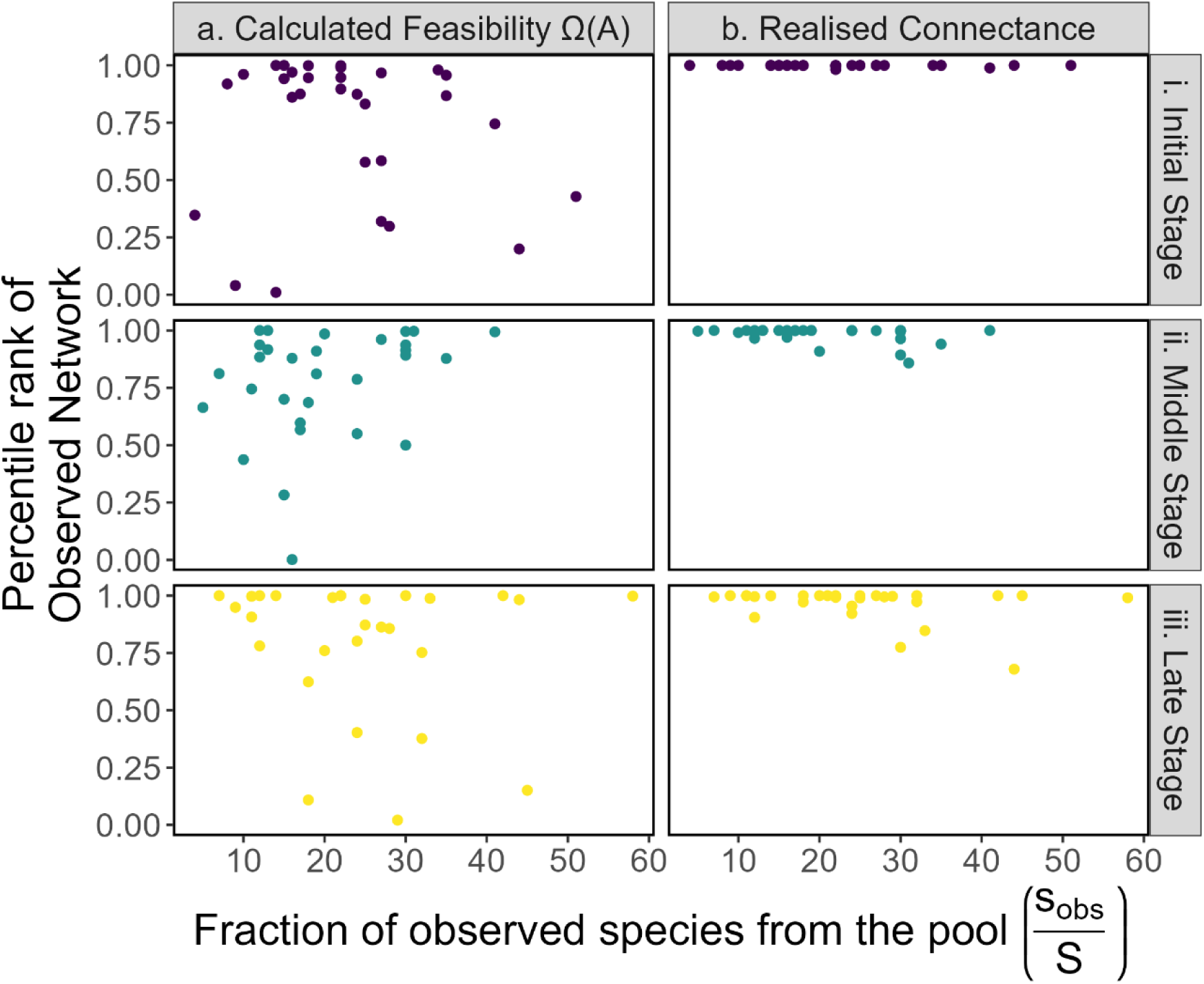
Comparison of properties (calculated feasibility domain and connectance) of observed tropical forest networks to randomly assembled communities. While estimated feasibility was significantly higher than would be expected under a random scenario (a), the signal from simple connectance was much stronger (b). This held true across a wide range of proportion of observed species (x-axis). Split of the communities into three stages and on the x-axis by the relative diversity of possible species seen in each community follows the original paper.

### Revaluating the relationship between community dynamics, **Ω** and network structure

The bi-directional relationship between community structure and dynamics has a long and complex history, bedevilled by the multitude of measures of both (Dunne & Pascual 2006). Given the assumptions underlying the methodology and uncertainties in the underlying estimated interaction matrices, calculated empirical values of Ω are unlikely to actually represent the theoretical quantities they purport to, yet might still have predictive power. This has potential consequences for the long-running discussion regarding the relationship between dynamics, feasibility and network structure (Aparicio *et al*. 2023; Bastolla *et al*. 2009; Blüthgen & Staab 2024; James *et al*. 2012; Rohr *et al*. 2014; Saavedra *et al*. 2016; Saavedra & Stouffer 2013; Suweis *et al*. 2013; Valdovinos *et al*. 2016). Rather than feasibility explaining the apparent dominance of certain network structures (such as high generality or nestedness), these results suggest causation appears as, if not more, likely to be the other way – the inferred Ω values of observed networks are higher because something about their underlying structure leads to higher calculated Ω values and higher likelihood of being observed as a community (Fig. 8). For example, García-Callejas *et al*. (2023) showed how inferred feasibility is closely driven by intraguild interaction overlap and average diagonal dominance across 18 communities. Observed networks have degree distributions that appear to maximise structural stability (Domínguez-Garcia *et al*. 2024). These relationships can account for how structural stability has been able to generate accurate predictions despite uncertainties in the input matrices and contexts where the fine detail of network structure is likely to play a subsidiary role in determining dynamics.

**Figure 8.**
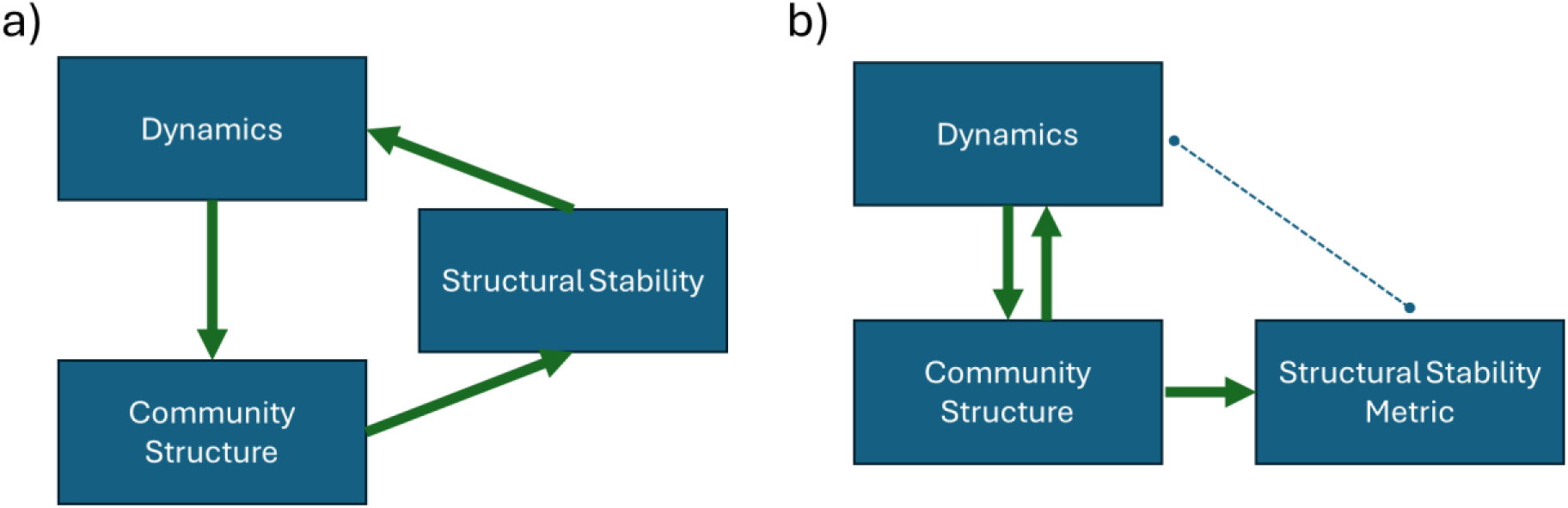
Alternative interpretations of observed correlations between structural stability and community assembly patterns. **a)** Idealised flow of information: structural stability synthesises something fundamental about the community structure that is a core driver of the dynamics of the ecological community. b) Possible relationships given empirical uncertainty and ecological complexity: the structural stability metrics that can be calculated will be driven by elements of community structure that could influence dynamics through some other mechanism, resulting in correlation without causation. In this scenario structural stability estimates would be less well correlated with the dynamics of interest than other calculable properties of community structure.

Increasingly, feasibility domain measures such as ω have been used as a stand-in response variable for describing the dynamic impact of network structure when actual dynamic data is not available (Arroyo-Correa *et al*. 2023; De Laender *et al*. 2023; Dougoud *et al*. 2018; García-Callejas *et al*. 2023; Rohr *et al*. 2014; Wu *et al*. 2025), a use that has been justified by its the apparent explanatory power in empirical tests. However, the weaker relationship to observed dynamics than simpler, more direct, metrics suggests that in empirical scenarios ω is often better thought of as a (noisy) descriptor of network ‘structure’, rather than an emergent dynamic response. Identifying the direction of causation between measures and importance is an important question - structural considerations such as feasibility may well be part of the deeper explanation for why connectance based metrics are able to perform well. It appears that ω is frequently a relatively inefficient summary of other network properties, and an overreliance of it as a response variable it may well distract from features, like basic connectance, that may be considerably more interpretable.

## Conclusion and Outlook

Structural stability and the analysis of feasibility is an inherently attractive approach, linking scales and ecological processes while apparently needing minimal information and unlocking a set of sophisticated analysis tools. It is crucial to stress that these case studies are not at all a rejection of the concept of feasibility as a theoretical tool for generating fundamental explanations for the patterns we observe in nature. Rather, they suggest that the precision required to apply the theory for real systems appears likely to be effectively empirically unattainable with current tools and datasets. Consequently, apparent successes need to be interpreted with more scepticism. If methods of generating interaction matrices from bipartite observations of interactions are effectively repackaging that qualitative structure in a pseudo-quantitative form, a key future question for future research is what can the calculation of feasibility add on top of other metrics?

It is well understood that the Lotka-Volterra assumptions necessary to calculate structural stability exclude numerous important processes that operate at the community level (Levine *et al*. 2017; Levins 1968). When studying complex, ever-changing, ecological communities, there will always be a fundamental gulf between the simplifications necessary to generate analytic results and the known complexity of real ecological systems (May 1973). Better connecting theoretical and empirical results must remain a driving objective, but the difficulty of the challenge means that the identification a correlation between abstract theoretical results and empirical observations has to be the start, not the end, of the road of identifying the utility of a framework.

The challenges of developing and testing theories around the role of species interactions in community assembly is well-trodden ground (e.g. Quinn & Dunham 1983; Roughgarden 1983; Simberloff 1983). Given the multi-causal, integrative, nature of ecology (Hilborn & Stearns 1982; Siegel & Dee 2025), explanations and contributing factors are often tightly interwoven and need to be carefully unpicked to form a firm basis for further work. This of course, is much easier said than done (Betini *et al*. 2017) but a key step to identifying if the assumptions and simplification are inherent within a particular framework are useful is comparing the explanatory power of an approach to alternative, more parsimonious, explanations. Until such a signal is demonstrated, the utility of assuming measures of Ω equate directly to the ‘likelihood of community persistence’, without further qualifiers, may be doubtful.

Moving forward, more explicitly accounting for parameter uncertainty in calculations is a basic start and can be directly propagated through to end results when underlying parameters are estimated within a Bayesian framework (Barabás *et al*. 2014; García-Callejas *et al*. 2023; Simmonds *et al*. 2022; Terry & Armitage 2024). However, such approaches still hit the fundamental barriers discussed earlier and may well not be enough to head off uncertainties inherent within the structural approach (Cervantes-Loreto *et al*. 2023). Analyses need to be clear exactly which contributors to uncertainty (model, parameters, environment, interaction structure, feasibility calculation etc.) are being propagated and how far through the analysis such uncertainty is carried through to ensure appropriate interpretation (Simmonds *et al*. 2024).

Beyond uncertainty propagation, comparative statistical approaches such as causal inference (Siegel & Dee 2025) can start to unpick the cause and effect of interrelated measures of community structure and dynamics, but will be restricted by the small effective sample size of comparable networks in most empirical studies. Shifts in the scale of communities analysed could also be productive - analyses focussing on smaller compartments of communities can allow higher accuracy with less scope for cascading dependences, but this cannot address dynamics of more complex communities. For large communities without very strong structure that meet stability criteria, feasibility is well determined by macro-properties of the community (such as connectance, mean interaction strength and variance, Grilli *et al*. 2017). By moving beyond the need for exact specifications of the interaction network, at larger scales there is scope to greatly reduce problematic over-sensitivity (Barbier *et al*. 2018).

Ultimately, the result that simple measures that don’t require detailed quantification of the whole network can better explain observed community assembly patterns can be interpreted as positive news for addressing the challenges of a changing world that ecologists hope to address (Dansereau *et al*. 2025). There is a rich and wide body of work that relates generality of species interspecific interactions to community dynamics at different scales (Armbruster 2017; Balisi *et al*. 2018; Carscadden *et al*. 2020; Colles *et al*. 2009; Dunne *et al*. 2002; Laurance 1991; Valdovinos *et al*. 2016). While developing more synthetic and wider-scope quantitative network level metrics has enormous value, local, species-level, measures that are more robust to uncertainty should not be neglected as a base from which to build a more holistic understanding of ecological dynamics.

## Acknowledgements

JCDT was supported by the Leverhulme Trust (ECF-2022-666). Constructive comments from Jie Deng, Chuliang Song, Alfonso Allen-Perkins & Virginia Domínguez-García on earlier drafts clarified some issues and discussions with the CERO and SalGo research groups helped consolidate the presentation.

