## Supplementary material for "Unfeasible expectations: the sensitivity of structural stability measures favours simpler metrics for empirical questions": SI

Supplementary Information

*Unfeasible expectations: why simple predictors outperform structural stability measures for understanding community assembly* Terry (2025)

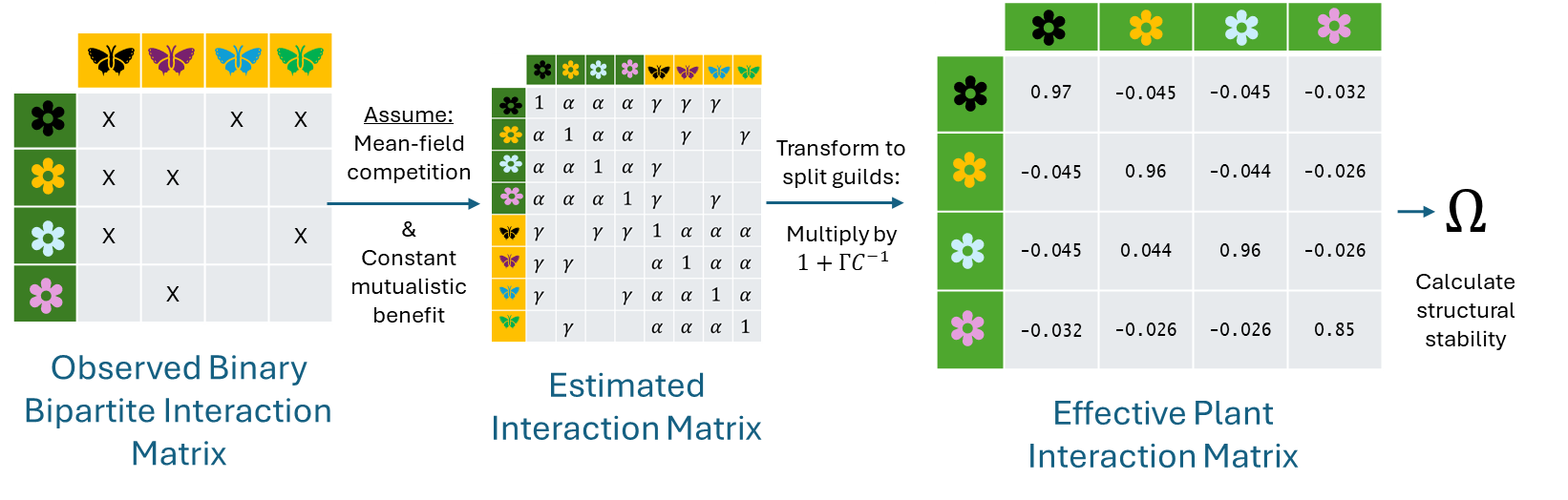

**Figure S1. Illustration of calculation of effective interaction matrix from niche overlap in binary bipartite networks used in case studies 1, 2, & 4 of the main text.** Taking as input a binary bipartite network of interactions (here pollination), a monopartite (united) matrix is inferred based on assumptions of mean-field competition within clades ($\alpha$) and mutualistic benefits ($\gamma$). The effective intraguild interaction matrix just between one clade is then extracted by diagonalising for each block, and then splitting out just the relevant section (see e.g. Domínguez-Garcia *et al.* 2024; Saavedra *et al.* 2016). Here results are shown using $\alpha$ = 0.005 and $\gamma_{ij} =\gamma_{0}{M_{ij}}/{k_{i}}$, where $k_{i}$is the degree of species $i$, $M_{ij}$ denotes a mutualistic link, and $\gamma_{0}$ = 0.1.

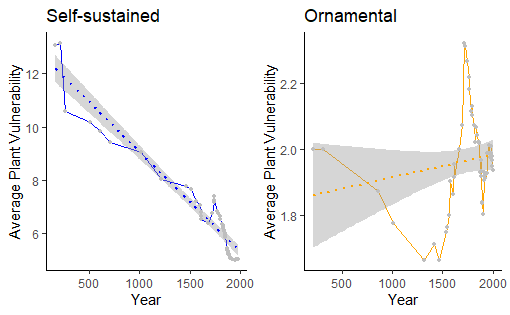

**Figure S2. Reproduction of the distinction between patterns of vulnerability and year for self-sustained and ornamental plants.** In the original paper, as a further piece of evidence supporting the value of feasibility in this scenario Song *et al.* (2018) show that $\omega$ could distinguish the assembly patterns of ornamental and non-ornamental plants, while a suite of standard network metrics (connectance of inferred competition network, NODF nestedness of plant-herbivore matrix, modularity of inferred competition matrices, mean competition strength of inferred competition matrices), showed the same trend in both ornamental and non-ornamental. However, repeating this test with vulnerability identifies an even stronger distinction between the two groups of species.  **Supplementary Text 1**: Boundary issues for the calculation of $\boldsymbol{\Omega}$ under uncertainty

The region of the feasibility domain can represent either locally dynamically stable coexistence if the intraspecific competition is stronger than interspecific competition or a dynamically unstable state (also termed contingent exclusion or priority effects, Song *et al.* 2021) in the reverse case when intraspecific competition is weaker than interspecific competition. In many theoretical analyses this challenge can be sidestepped by choosing values that guarantee stability, for example, the strong negative definite stability condition used by Grilli *et al.* (2017). Empirical analyses can also define their interaction matrices to ensure local stability (Saavedra *et al.* 2016). When the interaction matrices vary enough to cross this boundary, it is not immediately clear whether unstable domains should ‘count’. This categorisation challenge has been handled in three alternative ways for the two-species case where analytic formulae can be established:

1) To count either stable or unstable feasibility outcomes as part of a possible feasibility domain, effectively taking the absolute value of the difference between the boundaries of the feasibility domain. This is the approach taken by the multi-variate normal function used in the majority of multi-species studies, which does not test for the stability properties of the identified feasibility domain. An example pairwise analytic formula (Song *et al.* 2020a) is:

$$\Omega_{S20}=\frac{2}{\pi}arccos\left( \frac{\frac{a_{21}}{a_{11}}+\frac{a_{12}}{a_{22}}}{\sqrt{1+{\frac{a_{21}}{a_{11}}}^{2}}\sqrt{1+{\frac{a_{12}}{a_{22}}}^{2}}} \right)$$

2) To just count the stable feasibility domains, assigning the feasibility domain of unstable equilibria a size of 0. This is the approach taken by (Barabás *et al.* 2016) with their analytic formula:

$$\Omega_{B16}=\frac{2}{\pi}max\left[ \arctan\left( \frac{a_{22}}{a_{12}} \right)-\arctan\left( \frac{a_{21}}{a_{11}} \right),0 \right]$$

3) To measure the angle between the boundaries and assign the unstable domain a negative value. This is implication of formulae such as (Saavedra *et al.* 2017):

$$\Omega_{S17}=\frac{2}{\pi}arcsin\left( \frac{a_{11}a_{22}-a_{12}a_{21}}{\sqrt{a_{11}^{2}+a_{21}^{2}\sqrt{a_{12}^{2}+a_{22}^{2}}}} \right)$$

This discontinuity in interpretation has consequences when handling uncertain interaction coefficients and calculating an ‘average’ feasibility domain size from a posterior distribution of parameters. Examples using the posterior distribution of interaction coefficients at 24°C are illustrated in Fig.SI1.

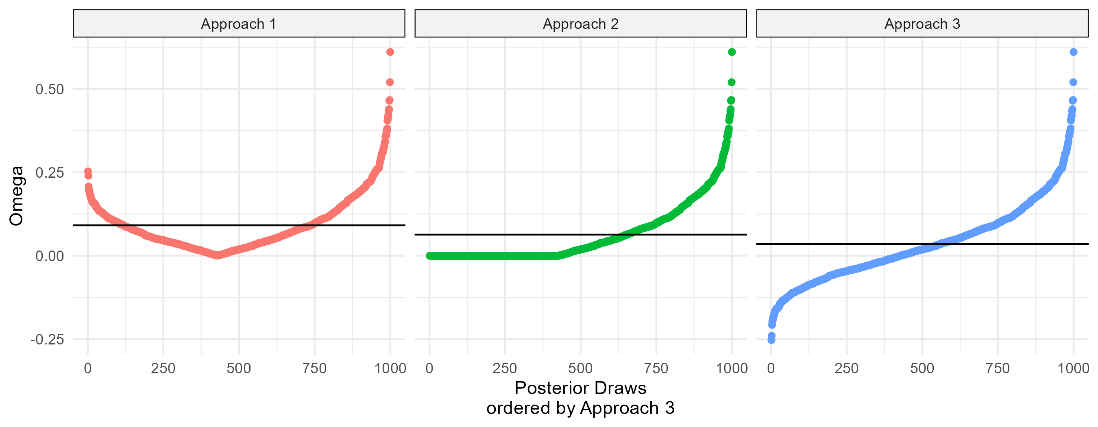

**Figure S3.** Example empirical posterior distribution of $\Omega$ values calculated by three analytical approaches used across the literature across 1000 posterior draws of competition coefficients at 24°C from the fitted models of Drosophila competition from (Terry 2025). Horizontal lines show mean values across the posterior. Note that the ‘average’ feasibility domain would be different in each case.

**Supplementary Text 2**: Model and parameters used in the empirical case study exploring uncertain parameters.

The population model fit in Terry (2025) describes the competition over a single generation in small vials between two *Drosophila* species at different temperatures: *D. pandora* (PAN) which favours warmer temperatures and *D. pallidifrons* (PAL) which favours cooler environments. The fitted model, determined after model selection, modelled the number of females and included an underlying thermal performance curve and temperature dependent Beverton-Holt competition:

$$N_{PAL,t+1}\sim N_{PAL,t}\frac{a_{PAL}\left( T_{t,}-T_{min, PAL} \right)\sqrt{T_{max, PAL}-T_{t}}}{1+\left( \alpha_{0PAL,PAL}+\alpha_{TPAL,PAL}T_{t} \right){(N}_{PAL,t}-1)+\alpha_{PAL,PAN}N_{PAN,t}}$$

and

$$N_{PAN,t+1}\sim N_{PAN,t}\frac{a_{PAN}\left( T_{t}-T_{min,PAN} \right)\sqrt{T_{max,PAN}-T_{t}}}{1+\left( \alpha_{0PAN,PAL}+\alpha_{TPAN,PAL}T_{t} \right)N_{PAL,t}+\alpha_{PAN,PAN}{(N}_{PAN,t}-1)}$$

where $a$, $T_{min}$ and $T_{max}$ are parameters of the thermal performance function defining the low-density growth rate, $\alpha_{ij}$ terms represent constant competition coefficients, whereas $\alpha_{0ij}$ is an intercept and $\alpha_{Tij}$ a slope term to linearly link a competition coefficient to temperature. Model errors followed a zero-inflated negative binomial. For PAL this included a temperature dependent zero-inflation term while for PAN the shape-term depended on the competition.

These models were fit in a Bayesian framework using the *brms* interface to STAN using a total of 1809 transition observations (1217 for PAL, 592 for PAN) assuming no observation error from the censuses. The fitted model is available in the code repository. From this core model, overall population growth rate parameters and competition coefficients were extracted for each temperature – the posterior of these composite parameters are shown in Fig S4:

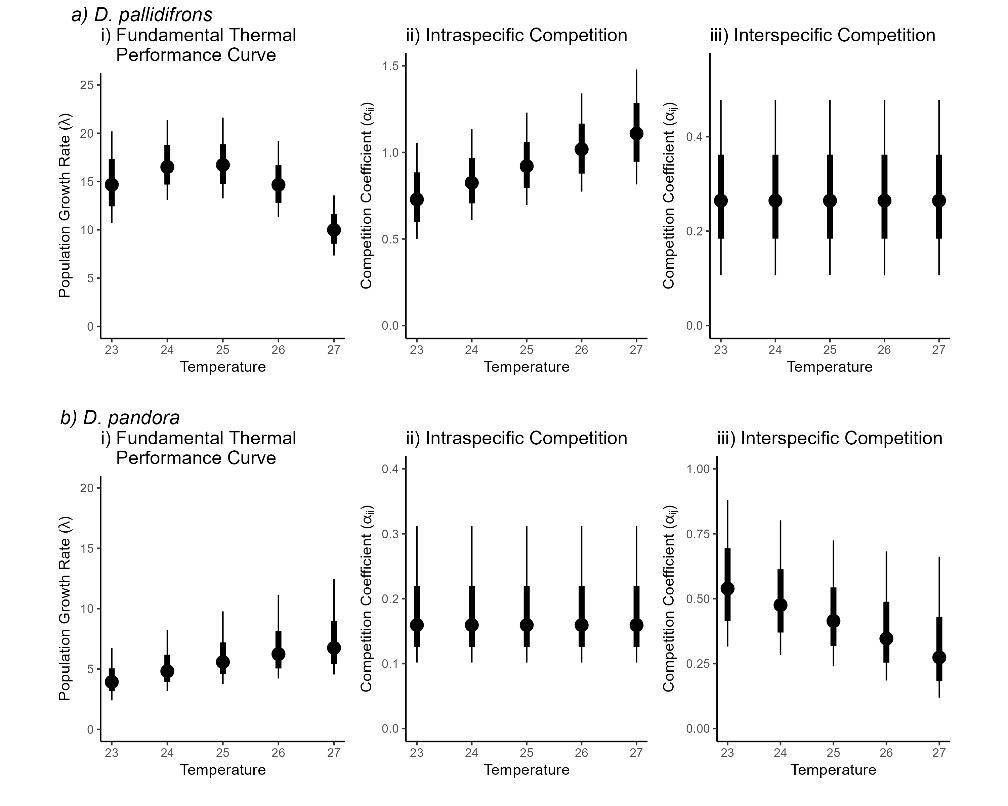

**Figure S4. Posterior parameter estimates across the five temperatures used in the main text figure.** Central value are medians, thick inner line is the central 66% percentiles, thin line is central 95% percentile.
